# Precision of auditory responses deteriorates on the way to frontal cortical areas

**DOI:** 10.1101/688549

**Authors:** Luciana López-Jury, Adrian Mannel, Francisco Garcia-Rosales, Julio C. Hechavarria

## Abstract

Frontal areas of the mammalian cortex are thought to be important for cognitive control and complex behaviour. These areas have been studied mostly in humans, non-human primates and rodents. In this article, we present a quantitative characterization of response properties of a frontal auditory area responsive to sound in the bat brain, the frontal auditory field (FAF). Bats are highly vocal animals and they constitute an important experimental model for studying the auditory system. At present, little is known about neuronal sound processing in the bat FAF. We combined electrophysiology experiments and computational simulations to compare the response properties of auditory neurons found in the bat FAF and auditory cortex (AC) to simple sounds (pure tones). Anatomical studies have shown that the latter provide feedforward inputs to the former. Our results show that bat FAF neurons are responsive to sounds, however, when compared to AC neurons, they presented sparser, less precise spiking and longer-lasting responses. Based on the results of an integrate-and-fire neuronal model, we speculate that slow, low-threshold, synaptic dynamics could contribute to the changes in activity pattern that occur as information travels through cortico-cortical projections from the AC to the FAF.

## Introduction

The prefrontal cortex is defined as a region of the anterior pole of the mammalian brain and it has been mostly studied in the brain of human and non-human primates (Goldman-Rakic, 1995). This area has been described as a highly integrative area dedicated to the representation and execution of goal-directed behaviours (Fuster, 2001). There is a wealth of neurophysiological studies that confirm its role in working memory, planning and decision making, using visual (Mikami *et al.*, 1982; Funahashi *et al.*, 1993; Tanibuchi & Goldman-Rakic, 2003; Murray & Rudebeck, 2018) as well as auditory paradigms (Plakke *et al.*, 2013; Plakke & Romanski, 2014).

In the last 20 years, the interest in auditory processing in prefrontal areas has increased (Medalla & Barbas, 2014). Several areas of the frontal lobe receive afferents from auditory processing regions (Hackett *et al.*, 1999; Romanski *et al.*, 1999a; Romanski *et al.*, 1999b), and studies have shown that neuronal responses reflect these projections since many prefrontal neurons are responsive to acoustic stimuli (Newman & Lindsley, 1976; Azuma & Suzuki, 1984). In addition to coding sensory stimuli, the prefrontal cortex is commonly regarded as a key area in cortical audio-motor integration (Fuster, 2000). Therefore, this area is involved not only in auditory cognition, multisensory integration (Watanabe, 1992; Rao *et al.*, 1997) and auditory attention, but also in cognitive control of the (vocal) motor system (Carmichael & Price, 1995).

Studies on non-primate animal models have postulated that prefrontal areas (as a functional block) are not unique to primates, although this idea continues to be controversial (Uylings *et al.*, 2003; Wise, 2008). In bats, a highly used animal model for auditory studies, a frontal area responsive to sounds has also been described (Eiermann & Esser, 2000; Kanwal *et al.*, 2000). This area was defined as the “frontal auditory field” (FAF). Due to their nocturnal habits and their ability to echolocate, bats rely strongly on the integration of complex auditory stimuli and on the motor coordination in relation to these sensory inputs, both in echolocation and communication contexts. It has been hypothesized that, much like prefrontal areas of primates, the bat FAF could be instrumental for the integration of sensory and motor information (Kobler *et al.*, 1987; Esser, 2003).

Interestingly, although the bat FAF was described more than 30 years ago, only a few studies dealing with this brain region have been published. Already in 1987, Kobler and colleagues showed, by means of anterograde tracing, a projection from the auditory cortex (AC) to the rostral part of the neocortex (the FAF) in the mustached bat (*Pteronotus parnellii*) brain (Kobler *et al.*, 1987). This area receives input from a second afferent pathway, which bypasses the inferior colliculus and projects directly from the medulla to the frontal cortex via the suprageniculate nucleus (Casseday *et al.*, 1989). It was hypothesized that the existence of this fast-afferent pathway hints towards the bat FAF as similar structure to the prefrontal cortex described in other mammalian species (Goldman-Rakic & Porrino, 1985; Kobler *et al.*, 1987). More than ten years after the first description of the bat FAF, two papers were published describing basic properties of FAF neurons in the mustached and the short-tailed fruit bat (*Carollia perspicillata*), in response to auditory stimuli (Eiermann & Esser, 2000; Kanwal et al., 2000). They reported recordings completely unlike the responses of the bat auditory cortical units. Unfortunately, after the FAF-related publications mentioned in the preceding text, no more studies have come to light regarding this structure, even though the need of studying frontal areas in bats has been discussed several times (Esser, 2003; Kanwal & Rauschecker, 2007; Kossl *et al.*, 2014).

The main aim of this study is to compare response properties of auditory neurons found in the bat FAF and AC. We wanted to assess if the quality of neuronal responses, quantified by measurements of response precision, latency, inter-trial variability (among others), changes between these two structures. For that purpose, using identical stimulation paradigms we performed electrophysiological recordings in the FAF and AC of adult *Carollia perspicillata* bats. Our results showed that FAF neurons are responsive to sounds but display a strong deterioration of response quality when compared to their counterparts in the AC. Based on the results of an integrate-and-fire neuronal model, we speculate that slow, low-threshold synaptic dynamics could contribute to the changes in activity patterns that occur as information travels through cortico-cortical projections from the AC to the FAF.

## Methods

The experimental animals (five males and two females, species: *C. perspicillata*) were taken from a breeding colony in the Institute for Cell Biology and Neuroscience at the Goethe University in Frankfurt am Main, Germany. The room had day/night – cycle (12 hours each) and the animals were fed daily ad libitum. All experiments were conducted in accordance to the Declaration of Helsinki and local regulations in the state of Hessen (Experimental permit #FU1126, Regierungspräsidium Darmstadt).

### Surgical procedures

The bats were anesthetized with a mixture of ketamine (100mg/ml Ketavet, Pfizer) and xylazine hydrochloride (23.32 mg/ml Rompun, Bayer). Under deep anesthesia, the dorsal surface of the skull was exposed with a longitudinal midline incision in the skin. The underlying muscles were retracted from the incision along the midline. A custom-made metal rod was glued to the skull using dental cement to fix the head during recordings. The animals then recovered at least one day from surgery. On the first day of the recordings, a small hole was made in the skull using a scalpel blade on the left side of the cortex in the position corresponding to either the AC or the FAF using the patterns of the blood vessels as a landmark (Esser & Eiermann, 1999; Hagemann *et al.*, 2010).

### Neuronal recordings

In all 7 bats recordings were performed over a maximum of 14 days with at least one day recovery time between each recording session. During the experiments the animals were always kept anaesthetized with a dose of 0.01 ml of the anesthetic mixture used for surgery (anesthesia was injected subcutaneously every 90 – 120 minutes). Each recording session lasted a maximum of 4h. The animals were positioned over a warming pad whose temperature was set to 27°C.

All experiments were performed in an electrically isolated, sound-proofed chamber. For the recordings, glass electrodes were pulled out of micropipettes (GB120F-10) with a glass puller (P-97 Flaming/Brown type micropipette puller; Sutter instruments). The electrodes were filled with a solution of potassium chloride (3 mol/l) and were fixed into and electrode holder connecting the electrode with a custom-made preamplifier with 10-fold amplification through a silver wire. Electrode impedance ranged from 4-12 MΩ. Electrodes were moved into position within the cortex with a piezo manipulator (PM 10/1, Science Products GmbH, Hofheim, Germany). The average depth of recordings in the AC was 333 μm (SD = 130) and 368 μm (SD = 102) in the FAF.

Recorded electrophysiological signals were filtered between 300 and 3000 Hz and amplified with a dual channel filter (SR 650, Stanford research). Signals were digitized with an acquisition board (DAP 840; Microstar Laboratories; sampling rate = 31.25 kHz) and stored on a computer for later analysis.

### Acoustic stimulation

All acoustic stimuli were synthetized using a custom-written Delphi software, generated by a D/A board (DAP 840; Microstar Laboratories; sampling rate = 278,8 kHz), attenuated (PA5, Tucker Davis Technologies), and amplified (Rotel power amplifier, RB-850). Sounds were then produced by a calibrated speaker (ScanSpeak Revelator R2904/700ß; Avisoft Bioacoustics, Berlin, Germany), which was placed 13 cm in front of the bats nose. The speaker’s calibration curve was obtained with a microphone (model 4135; Brüel & Kjaer).

We used 2 ms pure tones (0.2 ms rise/fall time) to characterize auditory responses. The sound pressure level of the pure tones was set to 60 dB SPL, and their frequency changed randomly in the range from 10-90 kHz (5 kHz steps, 50 trials).

### Neuron model

To model the spiking of FAF neurons, we used a simple leaky integrate-and-fire neuron model simulated with the Brian simulator (Goodman & Brette, 2009). In the model, the membrane potential (*V*_*m*_) is governed only by the subthreshold dynamic, after the following equation:

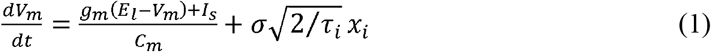

where *g*_*m*_ = 4 nS is the leak conductance, *E*_*l*_ = −50 mV is the resting potential, *C*_*m*_ = 1 pF is the membrane capacitance and *I*_*s*_ is the current from synaptic input. The neuron fires when *V*_*m*_ reaches the threshold *V*_*t*_ = −40 mV and then resets to *V*_*r*_ = −55 mV.

To allow for spontaneous activity, the neuron receives a random noise current added as an Ornstein-Uhlenbeck process with SD *σ* = 7 mV and time constant *τ*_*i*_ = 10 ms, given by the last term of the equation (1).

The equation defining the synaptic current,

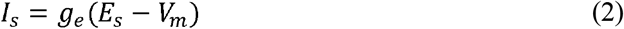

includes the synaptic reversal potential, *E*_*s*_ = 0 mV (standard for excitatory synapse), and the time varying synaptic conductance, *g*_*e*_, which evolves according to the differential equation

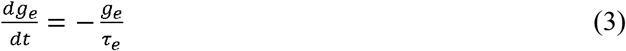

where the time constant *τ*_*e*_ determines the time course of the synaptic conductance change. When a spike arrives at a synapse, the synaptic conductance changes according to

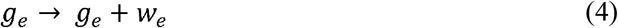

where *w*_*e*_ is the synaptic weight, a positive value of conductance. There is an intrinsic delay in the synaptic response dominated by the membrane time constant. In addition to the delay caused by membrane properties, in the model we also included an axonal delay of 15 ms.

All the parameters were set to reproduce the extracellular activity of FAF neurons. To ensure the latter, all values used were chosen so that they encompassed the range of values described for most of the neurons studied (see results). To study the effects of the synaptic dynamic on the spiking of FAF neurons, we performed several simulations changing two parameters: the magnitude of the synaptic weight, *w*_*e*_ and the time constant of the synaptic conductance change, *τ*_*e*_.

We simulated a single postsynaptic integrate-and-fire neuron, representing an FAF neuron, receiving an excitatory input through one synapse. The presynaptic spikes were generated from a Poisson process, in which the mean frequency of spike occurrence increases from 2 to 200 Hz for 25 ms after 10 ms of simulation, and then it returns to 2 Hz until the simulation ends. Note that a one-synapse simulation is a simplified situation in which it is assumed that the feed-forward input arriving to the FAF from the AC is strong enough for triggering spikes. We chose this approach because data from both the FAF and the AC obtained using identical stimulation paradigms were also available in this study. Electrophysiological data pertaining auditory responses in the *suprageniculate* nucleus (a structure that also provides input into the FAF (Kobler *et al.*, 1987; Casseday *et al.*, 1989)) are, to our knowledge, not available at present.

### Data and simulations analysis

To extract spikes waveform from the filtered signal, we selected a 4 ms time window around peaks whose amplitude was at least three standard deviations above the recording baseline. The waveform recorded on these time windows were sorted using a principal component analysis (PCA). For spike sorting we used an automatic clustering algorithm, “KlustaKwik”, that uses the obtained results from the PCA to create spike clusters. For further analysis, for each recording, we considered only the cluster with the highest number of spikes.

All the recording analysis were made using custom-written Matlab scripts (MATLAB R2018b, MathWorks, Natick, MA). Simulations and their analysis were done using Python. All the analysis relative to physiological data, except for the frequency tuning curve, were done considering those trials corresponding to the best frequency response of each neuron.

The estimation of post-stimulus time histograms (PSTH) from the real and simulated spike times were smoothed using a gaussian kernel function with a bin size of 1 ms and a bandwidth of 5 ms. As a measure of the precision (duration) of the response, we calculated the half width half height (HWHH) of the autocorrelogram of the smooth PSTH (sPSTH) as described above. The inter-spike interval distributions were also estimated using a kernel density estimator with a bin size of 1 ms and a bandwidth of 1 ms, in a range from 0 to 150 ms. The signal-to-noise ratio from frequency tuning curves was calculated as the maximal response (spike count) divided by the average of responses for all tested stimuli.

## Results

### Sparse, long-lasting spiking is a fingerprint of FAF responses

The FAF receives auditory afferents from the AC (Kobler *et al.*, 1987). In order to compare FAF and AC responses a total of 100 units, 50 from each area, were recorded extracellularly in response to pure tones at different frequencies (from 10 kHz to 90 kHz, 5 kHz steps; 60 dB SPL) in the short-tailed fruit bat *C. perspicillata*.

Like AC neurons, FAF neurons displayed evidence of frequency tuning. Examples of frequency tuning curves are shown in the first row of Figure 1, for an AC unit (Fig. 1A) and for an FAF unit (Fig. 1B). Both examples showed a preference for low frequency sounds. The response to tones, determined from the number of spikes fired between 10 and 150 ms after stimulus onset, was maximum at 15 kHz for the exemplary AC unit and at 20 kHz for the FAF one. Fig. 1 C-D show the raster plots and Fig. 1E, the post-stimulus time histograms (sPSTH, see Methods; smoothed used a 5 ms gaussian kernel, 1 ms resolution), obtained for both example units at their respective best frequency (BF). Simple visual inspection of the raster plots and sPSTHs already hints of very clear differences between FAF and AC responses in terms of their temporal response patterns: FAF responses to sounds are longer-lasting and less clear (in terms of number of spikes per time bin) than AC responses. Longer-lasting, broader sPSTHs rendered broader autocorrelation curves for the FAF example (Fig. 1F). In the autocorrelograms, we measured the half-width half-height (HWHH, gray horizontal line in Fig. 1F) as an indicator of response duration or temporal precision, as reported in previous studies (Kayser *et al.*, 2010; Garcia-Rosales *et al.*, 2018). The HWHH of the example FAF unit shown in Fig. 1F was 98.7 ms, more than double than that of the AC unit represented in the same figure (11.9 ms).

**Figure 1.**
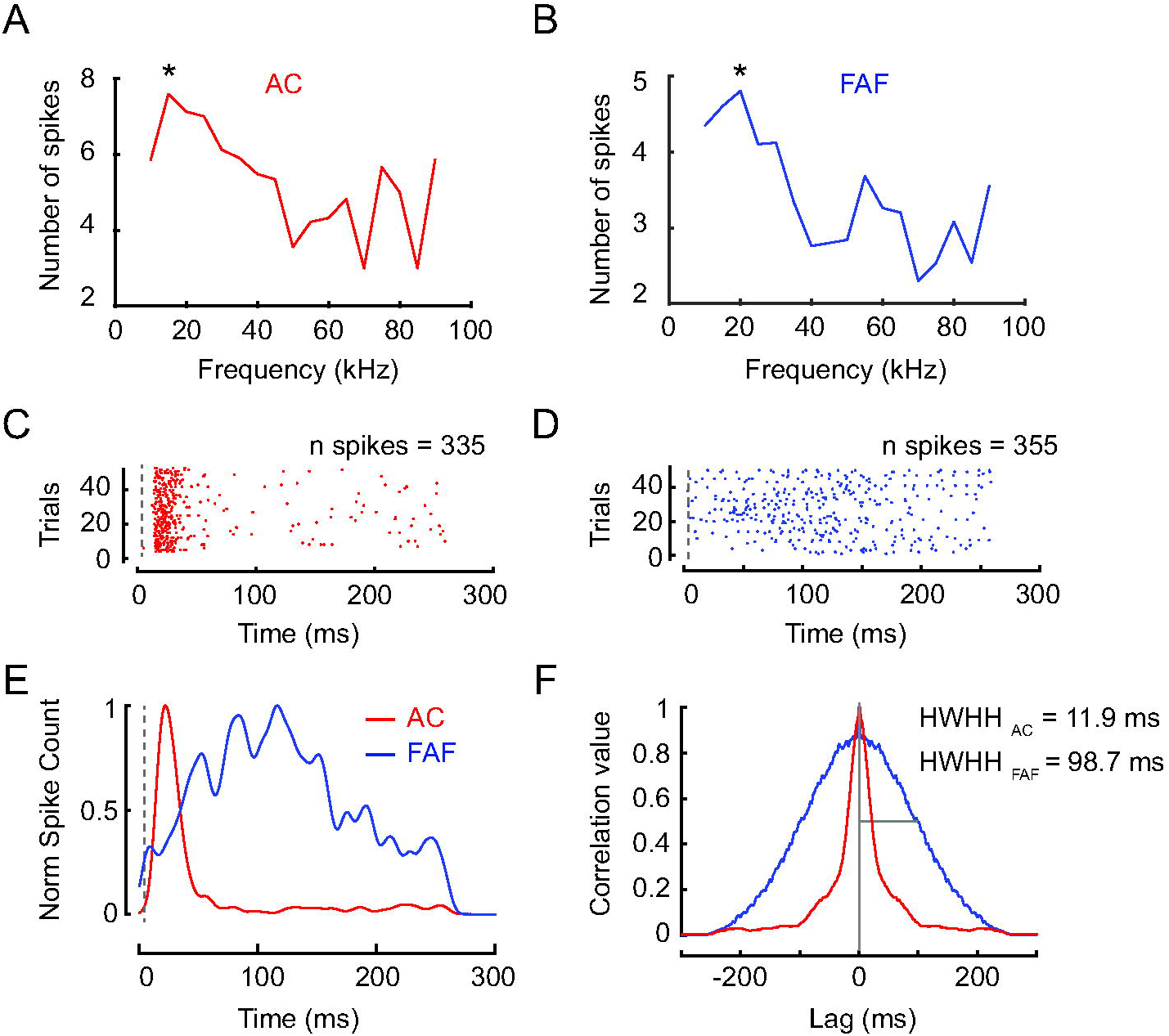
Two examples of low frequency tuned neurons recorded from the AC and the FAF. (A) and (B) show the number of spikes in response to pure tones at several frequencies for each unit. The number of spikes was measured as the mean of spike counts across 50 trials (from 10 and 150 ms after the pure tone onset). The asterisk indicates the BF and (C)-(F) plots correspond to the responses to these frequencies. (C) and (D) show raster plots and E, the corresponding sPSTHs for each neuron. The dashed line indicates the onset of stimulus. (F) shows the correlation values calculated from sPSTHs in the left. Grey lines represent the HWHH of the autocorrelation.

In order to establish if these differences were consistent across all recorded units, we made a population analysis statistically comparing each cortical area (Fig. 2). There were no significant differences between the population of FAF and AC units recorded regarding their BF distributions (Kolmogorov-Smirnov test, p-value=0.678, Fig. 2A). However, the two structures did differ according to the signal-to-noise ratio (SNR) of their tuning curves (Wilcoxon rank sum test, p-value=3.07*10^−7^, Fig. 2B), with the AC having higher SNR than the FAF. The SNR is an indicator of how much the response at the BF differentiates from responses to other frequencies and it is calculated as the spike count at the BF divided by the average of the spike count for all tested frequencies. Thus, higher SNR values indicate better tuning.

**Figure 2.**
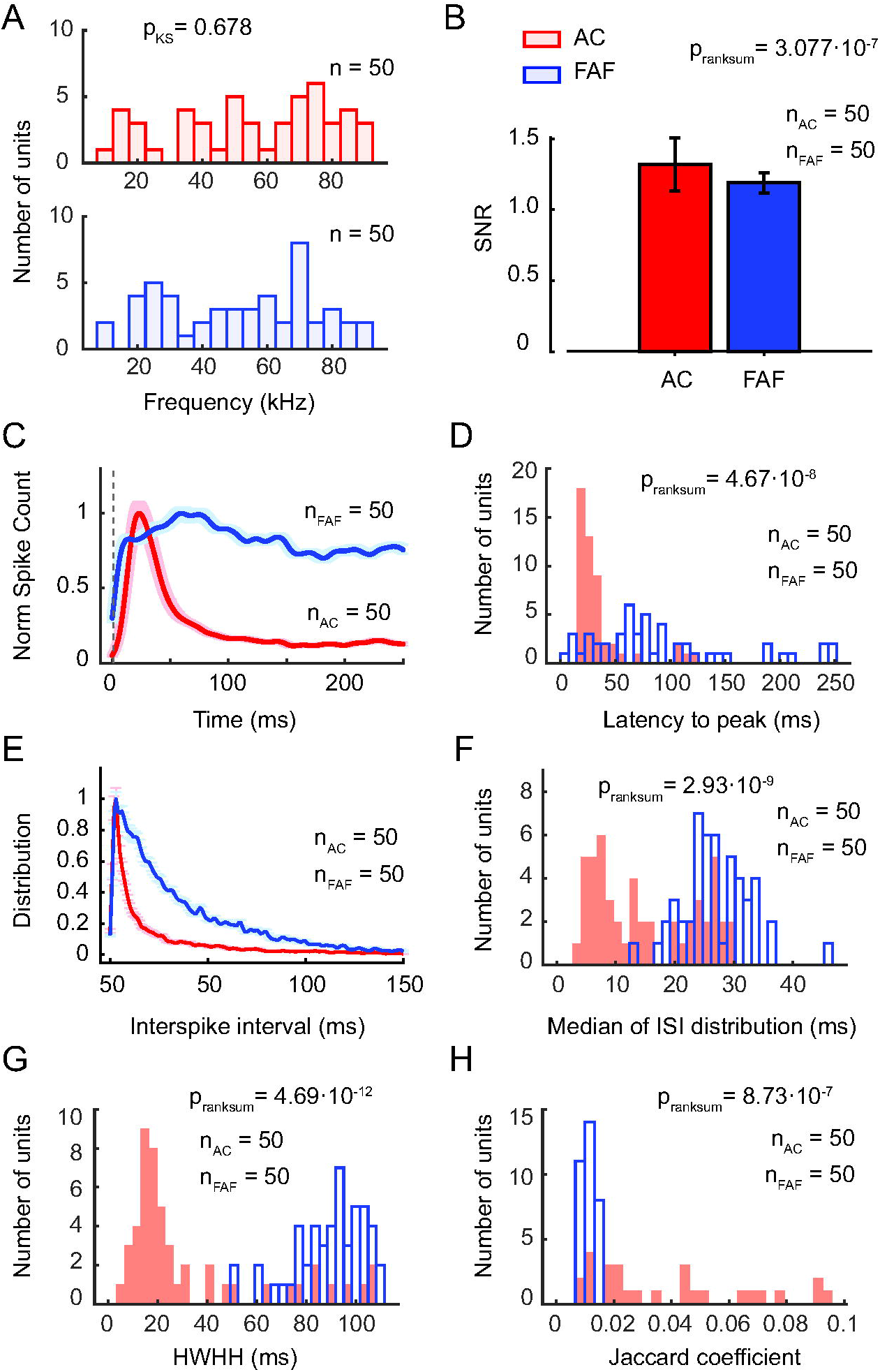
Population data comparing firing properties of 50 units recorded in the AC and 50 in the FAF. (A) shows the distributions of BF obtained in the AC (top) and in FAF (bottom). (B), the mean of signal-to-noise ratio (SNR) from frequency tuning curves, calculated as spike count at BF divided by the average spike count. Error bars indicate standard deviation. All the parameters analysed in (C)-(H) were based on the neuronal response to the BF. (C) shows the normalized average of sPSTH from all recorded units; shadow area indicates the standard error and the vertical dashed line, the onset of stimulus. (D), distributions of peak-response latencies calculated from sPSTH obtained from all units in AC and FAF. (E) shows normalized average of ISI distribution obtained in each location: AC and FAF. (F), distributions of median ISI for all units in AC and FAF. (G), distributions of HWHH obtained from the autocorrelograms of sPSTHs in each location. (H), distributions of averages of Jaccard coefficient calculated between trials as a measure of inter-trials variability, in AC and FAF.

Figure 2C shows the average sPSTH obtained at the BF for the units recorded in the FAF and AC. The curves were normalized to visualize the differences in the response time course in the two cortical areas studied. Average sPSTHs suggest that activation in the AC precedes responses in the FAF in most of the cases. To confirm the latter, we measured peak-response latencies in each neuron, calculated as the time between the onset of stimulus and the peak of the sPSTH. The distributions of peak-latencies for the two cortical areas studied are shown in Figure 2D. Peak-response latencies in the FAF ranged from 6 to 251 ms with an average of 93.4 ms (SD = 66). In contrast, AC units presented a narrower range of latencies with a lower average of 34.1 ms (SD = 25). The two distributions were significantly different from each other (Wilcoxon rank sum test, p-value=4.67*10^−8^). The percentage of units which present latencies lower than 20 ms was 10% in the FAF and 22% in the AC.

To study the spike-timing patterns, we calculated the inter-spike interval (ISI) distribution in all units and plotted the normalized average distribution per cortical area (Fig. 2E). The ISI distribution quantified for AC neurons was shifted to lower time intervals when compared to the FAF distribution. This result indicates that spiking in FAF neurons is slower than in the AC. The latter was also confirmed by analyzing the distribution of median ISIs calculated for each unit (Figure 2F). In the FAF, the average median ISI was 26.9±5.7 ms while in the AC that value amounted to 15±8.7 ms and the two ISI distributions were significantly different from each other (Wilcoxon rank sum test, p-value=2.93*10^−9^).

As mentioned in the preceding text, the duration/precision of responses can be calculated by means of the HWHH obtained from autocorrelograms of the sPSTHs (in each unit, only the sPSTH at the BF was considered, see Fig. 1). The histograms of HWHH obtained for the FAF and AC indicate that, in general, FAF neurons presented significantly longer response duration (and thus lower temporal precision) than AC neurons (Fig. 2G, Wilcoxon rank sum test, p-value=4.69*10^−12^). The mean HWHH obtained for the FAF (88.9 ms, SD = 14.6) was more than double that observed in the AC (33.9 ms, SD = 29.1) thus indicating that temporal spiking precision deteriorates on the way to frontal areas.

In the preceding text, we mentioned that FAF neurons had irregular discharge patterns. To quantify inter-trial variability, we used the Jaccard coefficient (see Methods). This metric was used in previous neurophysiological studies in bats (Macias *et al.*, 2016) and it allows to estimate the similarity between two binary words. Thus, we first transformed spike time series from each BF trial into a binary word (1= spike, 0=no spike, bin size = 1ms). Next, for each pair of trials, we extracted the Jaccard coefficient and then average the results obtained across all possible trial combinations in each neuron. The values can range from 0 to 1 (1 indicates equal binary words). Figure 2H shows that the response pattern similarity across trials is statistically lower in FAF than in AC neurons (Wilcoxon rank sum test, p-value=8.73*10^−7^). The latter adds to the idea that FAF neurons present more variable discharge patterns (in this case across trials, but also within trials, see preceding text).

Several studies have found that different types of neurons can be distinguished by the duration of their action potentials (Wilson *et al.*, 1994; Constantinidis & Goldman-Rakic, 2002). We examined whether spike widths differed between the FAF and AC to gain an insight into the spiking mechanisms in these two cortical areas. We hypothesized that the slow spiking dynamics of FAF neurons could be due to intrinsic membrane properties that could also influence spike shape (for review see (Bean, 2007)). This was not the case. Figure 3A shows the mean waveforms for 50 neurons recorded on each cortical area studied and the average of these means (the height of the waveforms was normalized to aid in comparing spike widths). Surprisingly, waveforms from the FAF and AC had similar shapes and durations. To quantify the shape of the spikes, we calculated the area under the absolute value of the average spike waveform for each unit studied. The area-under-waveform distributions (Fig. 3B) did not differ statistically between the AC and FAF (Wilcoxon rank sum test, p-value=0.357). The latter suggests that similar current dynamics are involved in spike generation in these two structures.

**Figure 3.**
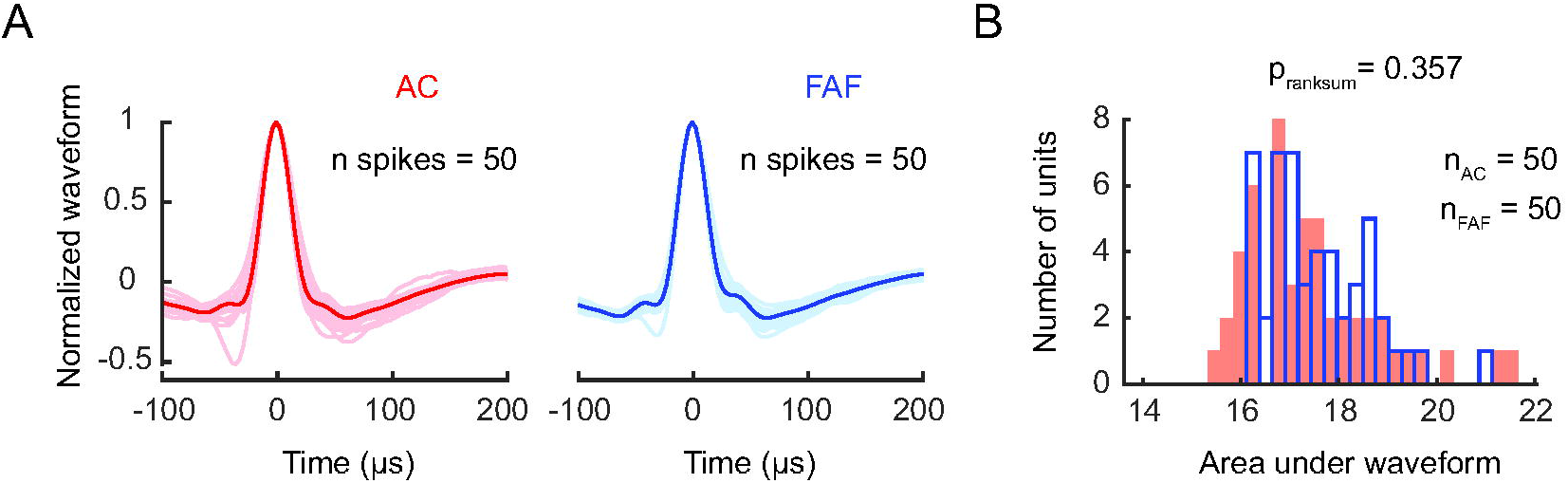
No differences in spike waveforms between units from AC and FAF. (A), average waveforms of each of the 50 units recorded in the AC (left) and in the FAF (right). The thicker line indicates the mean of the averages. (B), distributions of mean areas under the absolute value of the spike waveform obtained in each location (AC and FAF).

### Subthreshold, long-lasting excitation could explain FAF response properties

Our electrophysiological data showed that FAF neuronal properties differ strongly from those found in the AC. In comparison to AC neurons, FAF neurons have slower spiking (in terms of ISI) and longer lasting responses, which are loosely time-locked to auditory stimuli. It is unknown which mechanisms shape the firing properties of FAF neurons, causing responses in this structure to differ from those found in the AC (this last structure provides afferent information to the FAF). To address this issue, we created a simple leaky integrate-and-fire neuron model of an FAF neuron. The cellular properties of our model neuron (see Methods Eq. (1)) are specified by seven parameters. We constrained the parameters by comparing the simulated responses of the FAF model neurons to the experimental data.

We modeled the input to the FAF neuron model (that is, the response of AC projections driving FAF neurons) as a Poisson point process mimicking the average spiking of AC neurons. Figure 4A shows the raster plot from a simulation of 50 Poisson processes and the respective event counts below (sPSTH, bin size of 5 ms). Note that both the simulated and the observed AC (Fig. 2C) sPSTHs represent transient responses lasting less than 100 ms.

**Figure 4.**
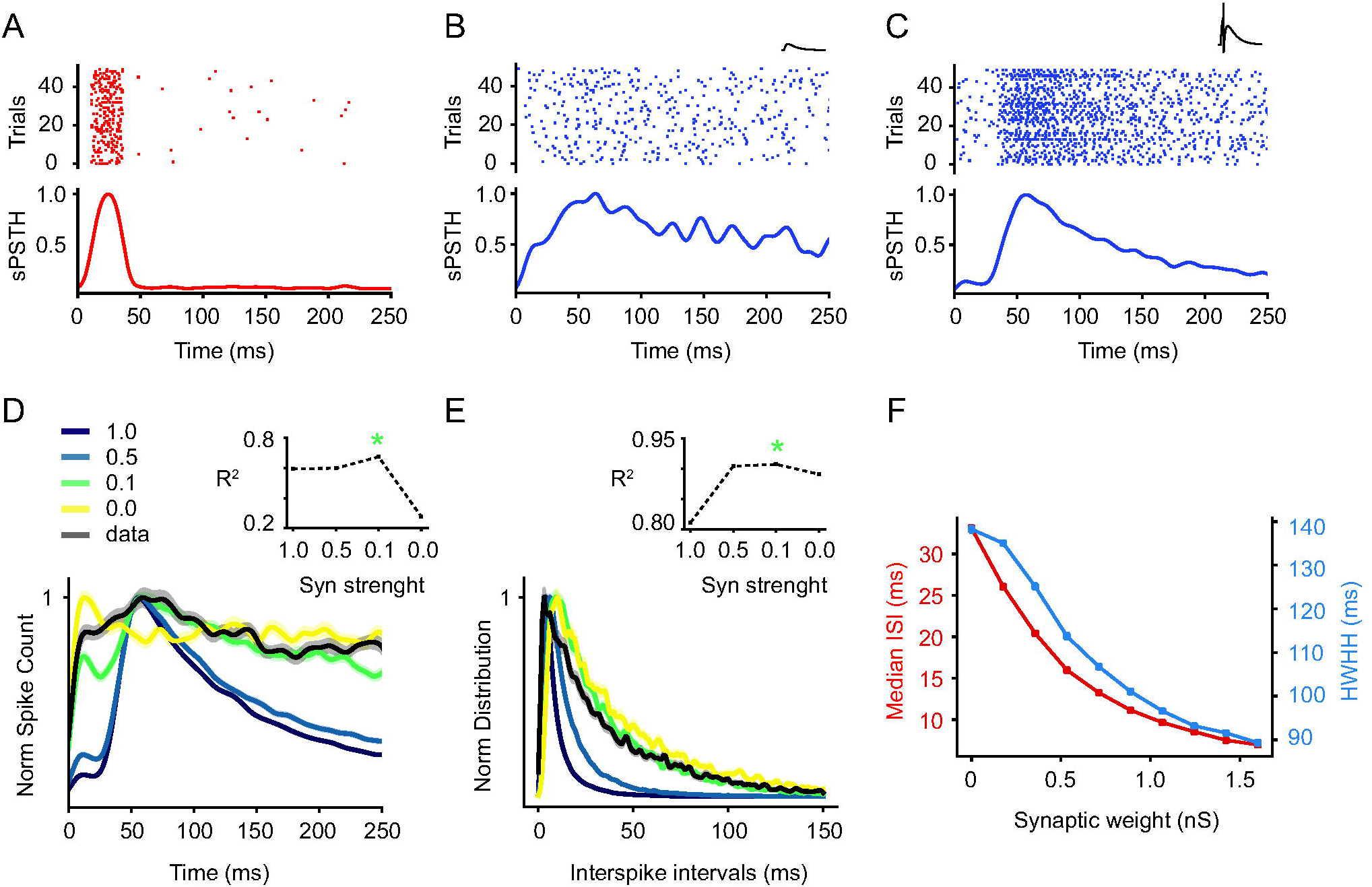
A simple model of FAF neuron suggest that subthreshold synapse could explain the observed spiking pattern. (A) shows a raster plot generated by a Poisson process mimicking one AC neuron response to a pure tone, below the corresponding sPSTH. (B) shows the spiking of 50 neuron models in response to the poissonic input shown in A. On the top, the raster plot and below, the corresponding sPSTH. The synaptic weight was set to 0.3 nS in order to generate a subthreshold excitatory post synaptic potential as indicated in the inset. The value used corresponds to a 18% of the minimum suprathreshold synaptic weight (1.6 nS). In (C), the same plots than in (B) are shown, however the synaptic weight in these simulations was set to 1.6 ns. One synapse is able to generate a spike in the neuron model, as indicated in the inset. In (B) and (C) the time constant of the synapse model was set to 90 ms. (D) shows sPSTH obtained from four simulations using different values of synaptic strength, defined as the ratio between the parameter *w*_*e*_ used and the minimum value of *w*_*e*_ necessary to reach a postsynaptic spike. The synaptic strengths used are shown in the legend. The average sPSTH from real data is plotted in black. The inset shows the Pearson correlation coefficient calculated between each simulated sPSTH and the average sPSTH obtained from all units. The asterisk indicates the higher value. (E) shows the normalized ISI distributions for the same simulations ran in (D). The inset shows the Pearson correlation coefficient calculated between these curves and the average ISI distribution obtained from all units. The asterisk indicates the higher value. (F) shows the median of ISI and the HWHH of autocorrelogram of sPSTH calculated from 10 simulations in which systematically increasing the strength of the synapse, from 0 to 1.6 nS. Error bars indicate standard errors across 50 simulations.

Since we showed that neurons in the AC and FAF exhibit statistically similar spike waveforms (see Fig. 3), we assumed that the modifications to input-output features of FAF neurons relay on synaptic properties more than on intrinsic neuronal properties. This assumption was based on several studies that have shown that the spike shape reflects kinetics and distribution of ion channels (Martina & Jonas, 1997; Henze *et al.*, 2000). Therefore, we investigated the effects of the synaptic weight and duration on the spiking of the FAF neuron model.

First, we investigated how the magnitude of the synaptic depolarization changed the firing of the FAF neuron model. Our results indicated that the postsynaptic effect must be subthreshold to reproduce the FAF data. In our model, a presynaptic spike induces conductance changes according to Eq. (3), (4) and (5) (see Methods section), which can produce (or not) a postsynaptic spike depending on the strength of the synapse. According to the set parameters, the minimum suprathreshold synaptic weight of our neuron model is 1.6 nS. Therefore, to generate a subthreshold excitatory postsynaptic potential, we set the synaptic weight at 0.3 nS, which corresponds to 18% of 1.6 nS. In Figure 4B we plotted the raster and the corresponding sPSTH from 50 FAF simulated neurons (or trials, no difference in our model), in response to the spike trains showed in 4A, when one presynaptic spike generates a small depolarization in the postsynaptic membrane potential as shown in the inset. The obtained sPSTH was qualitatively similar to the one obtained in FAF recordings (see sPSTH in Fig. 2C). In contrast, when the modeled AC-FAF synapse was strong enough to produce a postsynaptic spike (synaptic weight = 1.6 nS, see inset in Fig. 4C), the simulations rendered a sPSTH with high amplitude response, qualitatively different from those observed in our FAF data. In both simulations (Fig. 4B – C), the synaptic time constant was 90 ms (see below).

In order to systematically examine the effect of the synaptic strength on the shape of the sPSTH obtained from 50 neuron models, we ran four different simulations varying the synaptic weight (Fig. 4D) and compared them directly with the average data from all FAF units recorded (black curve). Here, we refer to synaptic strength as the ratio between the synaptic weight (*w*_*e*_, described in Eq. (4) in Methods) and the minimum *w*_*e*_ needed to generate one spike in our FAF neuron model:

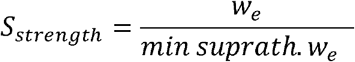

We tested synaptic strength values of 0, 0.1, 0.5 and 1. As shown in Fig. 4C, simulations with suprathreshold synaptic strengths (synaptic strength = 1) rendered an sPSTH with a short but high amplitude peak (result not shown) in response to the input, qualitatively different from that obtained in the data. When the synaptic strength was decreased to 0.5, the FAF simulated responses decreased in amplitude (result not shown) and increased in duration, however, the sPSTH differed from that obtained from FAF data. When the synaptic strength was set to 0.1, we obtained simulation results that resembled more the FAF data, consistent with a high value of Pearson correlation coefficient (r = 0.66) between simulated sPSTH and average sPSTH obtained from FAF recordings (indicated with an asterisk in the inset). As expected, when the synaptic strength was set to 0, there was no evoked response and the sPSTH reflected the spontaneous activity implemented in the neuron model.

In the preceding text, we also reported that FAF neurons have slow spiking dynamics (shown in higher values of median ISI when compared to the AC, see Fig. 2F). Therefore, for the same four simulations varying the *w*_*e*_, we calculated the ISI distributions (Fig. 4E). For decreasing values of synaptic strength, the ISI distribution shifted to higher time intervals, indicative of slower spiking. Again, setting the synaptic strength to 0.1 led to the highest Pearson correlation coefficient (r = 0.91, indicated with an asterisk in the inset) between the ISI data distribution and the four different simulation distributions.

To summarize these results, in Figure 4F, we ran 10 simulations covering a larger range of subthreshold synaptic strengths and plotted, for each simulation, the median ISI and the HWHH. Both parameters decrease with increasing the synaptic strength, indicating that stronger synapses reduce the differences between input and output spiking. Based on these simulation results, we hypothesize that the increased HWHH and ISI observed in the FAF when compared to the AC could be the result of weak excitability power in the AC-FAF projection. Note that with our model we cannot disentangle whether this weak excitability power is the result of weak synapses or low resting membrane potentials in FAF neurons. The latter could result, for example, from inhibitory regimes.

Next, we investigated how the duration of the synaptic depolarization changed the firing of the FAF neuron model. Figure 5A-B show simulations in which the time course of the synaptic conductance change, given by the parameter *τ*_*e*_, was set to 10 and 90 ms, respectively. The post-synaptic potentials modeled in each case can be seen in the respective insets. In both simulations, the synaptic strength was subthreshold and fixed to 0.1 (ratio compared to the minimum suprathreshold conductance, see preceding text). The simulated sPSTH showed that shorter excitatory post synaptic potentials (Fig. 5A) result in shorter responses that those obtained using longer time constants (Fig. 5B). Note that simulations with longer excitatory post synaptic potentials reproduced more closely the sPSTHs observed experimentally (see Fig. 2C).

**Figure 5.**
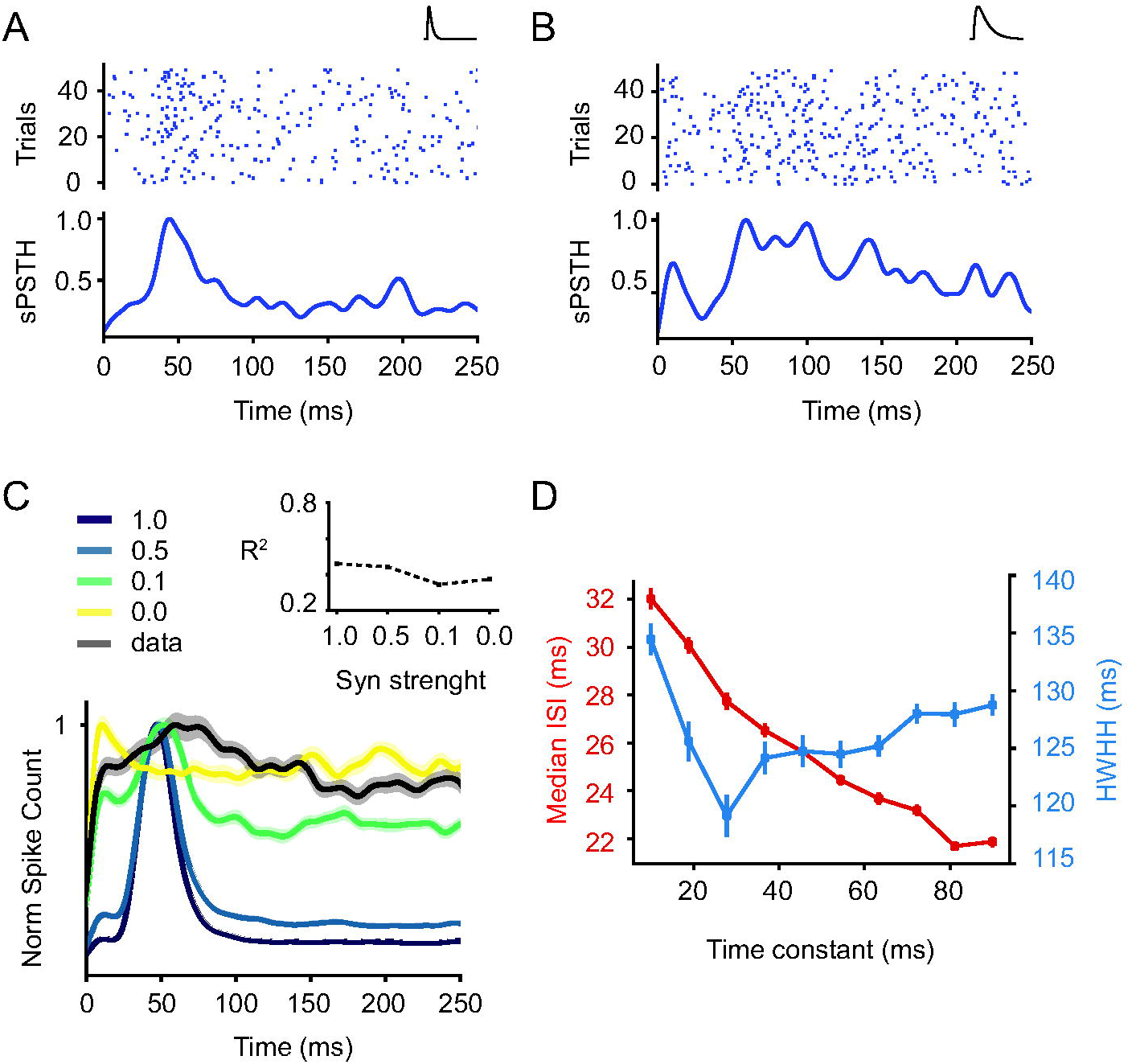
A simple model of FAF neuron suggest that long-lasting synapse is also needed to explain the observed spiking pattern. (A) and (B) show simulations in response to the spikes showed in Figure 4A, with different post synaptic durations. On top, the raster plot and below, the respective sPSTH. In (A), the time constant of the synaptic conductance change was set to 10 ms, the change in the membrane potential produced by one presynaptic spike is showed in the inset. In B, the time constant was set to 90 ms. Note that the excitatory post synaptic potential in the inset is longer in comparison with the inset in (A). In (A) and (B), the synaptic weight was set to 4.3 nS and 0.3 nS, respectively. Both values correspond to a 18% of the minimum suprathreshold synaptic weight of each model. (C) shows the sPSTHs obtained from four simulations changing the synaptic strength as showed in Figure 4D. The synaptic strengths used are shown in the legend. In these simulations, we set a shorter synaptic time constant than that in Figure 4D (*τ*_*e*_ = 10 ms instead of 90 ms). The average sPSTH from real data is plotted in black. Note that even with the lowest synaptic strength tested (ratio = 0.1), the Pearson correlation coefficient (showed in the inset) is lower than those obtained in Figure 4D, using longer synaptic time constant. (D) The median of ISI and HWHH of autocorrelograms of sPSTHs obtained from 10 simulations in which the duration of the postsynaptic effect was systematically increased, by changing *τ*_*e*_ from 10 to 90 ms. Error bars indicate standard error across 50 simulations.

To make sure that long synaptic conductance was indeed needed to reproduce experimental data, we ran the same simulations as in Figure 4C, however, in this case, the synaptic time constant was set to 10 ms (it was 90 ms in Fig. 4C). None of the simulations obtained using 10-ms time-constant were able to better reproduce qualitatively the experimental data (showed in black), consistent with lower values of Pearson correlation coefficients (r_max_ = 0.45) in relation to those obtained with 90-ms time-constant (compare with inset Fig. 4D). These R values (inset Fig 5C) were calculated by correlating the average data and the four different simulated sPSTHs. Overall the simulation results indicate that low synaptic strength is not enough to reproduce FAF responses. Long excitatory post synaptic potentials area also needed.

Finally, we ran 10 simulations in which the synaptic time constant was systematically increased from 10 to 90 ms. For each simulation, the median ISI and the HWHH from the neuron model output was analyzed (Figure 5D). The median ISI decreases with increasing time constant, indicating that longer excitatory post synaptic potentials enable an increment in the rate of temporal summation, and consequently the probability of spike occurrence increases. On the other hand, HWHHs changed in a complex U-shaped manner with increasing time constants. Note that the experimental data on FAF responses showed high ISI and HWHH values (Fig. 2). Based on our simulations, high ISI-HWHH combinations can be achieved only with long synaptic time constants.

## Discussion

The main aim of this article was to compare response properties of auditory neurons found in the bat FAF and AC regions. We used identical recording and stimulation settings for studying these two structures in order to assess modifications to neuronal responses in the AC-FAF pathway. Our results show that FAF neurons are responsive to sounds, however, when compared to auditory neurons, they presented sparser, less precise spiking and longer-lasting responses to pure tones. Based on results from a leaky integrate-and-fire neuronal model, we speculate that slow, weak synaptic dynamic could contribute to the changes in activity pattern that occur as information travels through cortico-cortical projections from the AC to the FAF.

### Possible origin and function of FAF responses

We found that, in general, peak response latencies to auditory stimuli are longer in FAF than in AC neurons. These latency differences suggest that cortical feed-forward projections evoke responses in the FAF. The latter falls in line with anatomical data showing that the FAF receives cortical afferents (Kobler *et al.*, 1987). However, this does not mean that AC inputs are the only source for driving FAF spiking. FAF neurons also receive afferents from auditory structures that bypass the AC, such as the suprageniculate nucleus of the thalamus (Kobler *et al.*, 1987). To our knowledge, detailed anatomical and electrophysiological data of suprageniculate afferents to the FAF are presently not available. How different inputs sources to the FAF (e.g. suprageniculate and cortical) interact at the single neuron level remains obscure. One could speculate that non-cortical inputs travelling through the suprageniculate nucleus could arrive to the FAF before cortical inputs. In our data, we did observe a small subpopulation of FAF neurons that had peak latencies shorter than 20 ms (10% of the recorded units, see Fig. 2D), although these neurons are not predominant. Similar results were described in previous studies on frontal responses in bats (Eiermann & Esser, 2000; Kanwal *et al.*, 2000). Note that with our data, we cannot disentangle whether neurons with fast latencies in the FAF result from fast projections from the AC or from fast afferents travelling through the suprageniculate nucleus.

Another possibility that requires consideration is that fast suprageniculate inputs could change the *status quo* in FAF neurons by controlling their membrane potential. Because most FAF neurons have long peak latencies, it is likely that suprageniculate afferents alone are not capable of evoking FAF spiking. The latter could occur either because suprageniculate inputs are excitatory but not strong enough for causing suprathreshold depolarizations, or because they are inhibitory. The inhibitory hypothesis is particularly appealing, as it could provide a likely explanation for achieving low FAF excitability which, according to our computational model, is a fundamental requisite for explaining FAF firing properties (se Fig. 4). In other species, GABAergic (inhibitory) projections from the Raphe nuclei and basal forebrain to frontal regions have been described (Carr & Sesack, 2000; Henny & Jones, 2008).

Our data show higher spiking variability (i.e. inter-trial variability calculated as Jaccard coefficients) in frontal neurons when compared to AC neurons (see individual examples in Fig. 1C and D and population data in Fig. 2H). This result is consistent with previous studies in bats and primates describing neuronal responses to auditory stimuli in frontal areas as irregular and variable (Newman & Lindsley, 1976; Eiermann & Esser, 2000; Kanwal *et al.*, 2000). Even though neural activity across sensory pathways increases in variability (Vogel *et al.*, 2005), studies indicate that this “variability” or output “noise” might offer processing advantages. Noisy responses could result from large excitability fluctuations in the neurons studied which could make these neurons more prone to control through processes such as attention or multi-sensory integration. In other animal species, these processes are known to influence frontal responses (Watanabe, 1992; Tomita *et al.*, 1999) and they could also do so in bats. Interestingly, studies have suggested that intrinsic noise could enhance sensitivity to weak signals (Stein *et al.*, 2005). The latter might be important for bats that have to cope with faint signals during navigation (i.e. echoes) often in noisy environments (Corcoran & Moss, 2017).

Note that it has been argued that response variability might be overestimated simply because we do not understand what high-order neurons are signaling (Masquelier, 2013). This could explain why responses from frontal neurons are less reliable than those of cortical neurons when tested with simple stimuli such as the pure tones used in this study. The reliability of neuronal responses could increase during active behavior (i.e. echolocation), which involves a set of variables such as attention, and the build-up of expectations based on previous sensory history (Feng & Ratnam, 2000; Wohlgemuth *et al.*, 2016). The roles of these variables is much diminished in experiments conducted in anesthetized, passively listening animals (this study). Future studies should investigate neuronal responses in awake animals during active behavior, to better understand auditory processing in frontal areas. In any case, it should be considered that experiments conducted under anesthesia could give insight into intrinsic properties of the neurons studied, disregarding how these properties might be influenced by complex phenomena such as attention.

### FAF response properties could be linked to the strength of afferent projections

To our knowledge, at present, histochemical, anatomical, and biophysical data regarding the bat FAF are very limited. We implemented a simple leaky integrate-and-fire neuronal model to gain insights into the cellular mechanisms that could explain the nature of FAF responses. Similar models have been widely used in neurophysiological studies for understanding sensory processing (Kremer *et al.*, 2011; Rossant *et al.*, 2011). Our aim with this model was to show, in the simplest configuration imaginable, how it is possible to achieve neuronal responses as those recorded in the bat FAF considering only AC feedforward inputs and their properties. Based on our empirical data, the FAF displays slow, irregular, long-lasting responses while AC spiking is reliable and temporally precise (see Fig. 2). Note that our model was based only on AC afferents. Extralemniscal inputs travelling through the suprageniculate nucleus could be even more reliable and precise than AC afferents (see preceding text).

Even though our model is very reductionist (for example, it does not consider either inhibitory inputs nor possible interactions within the FAF) it still was capable recreating basic properties of FAF neural activity based solely on long, weak excitatory post synaptic potentials (see Figs. 4 and 5). Both synaptic features suggest mechanisms by which FAF neurons can integrate different auditory activity over long periods of time by temporal or spatial summation of presynaptic inputs. Note that in our model we modified the “synaptic strength” to recreate FAF response properties. However, we cannot disentangle whether “synaptic strength” is linked directly to properties of pre-synaptic or post-synaptic neurons, or both. At the post-synaptic level, low synaptic strength could be caused, for example, by a low number of excitatory receptor channels or by very negative resting membrane potentials linked to inhibitory regimes (Wehr & Zador, 2003; Puig *et al.*, 2005). At the pre-synaptic level, low synaptic strength could be associated to a number of factors that ultimately decrease the amount of neurotransmitter that reaches post-synaptic neurons. In principle, all of the aforementioned mechanisms are plausible and not mutually exclusive ways to achieve low amplitude post-synaptic potentials in FAF neurons.

Our model also indicates that to achieve FAF-like responses slow depolarizations are needed (see Fig. 5). AMPA and NMDA receptors are the primary mediators of excitatory synaptic transmission in the cortex (Ozawa *et al.*, 1998). It is known that NMDA receptor-channels show slower kinetics than AMPA receptor-channels (Sanchez *et al.*, 2007). It is therefore tentative to suggest that NMDA could play a major role in mediating postsynaptic responses in FAF neurons. Future studies using pharmacological tools could test this prediction.

## References

Azuma, M. & Suzuki, H. (1984) Properties and distribution of auditory neurons in the dorsolateral prefrontal cortex of the alert monkey. Brain Res, 298, 343–346.

Bean, B.P. (2007) The action potential in mammalian central neurons. Nat Rev Neurosci, 8, 451–465.

Carmichael, S.T. & Price, J.L. (1995) Sensory and premotor connections of the orbital and medial prefrontal cortex of macaque monkeys. J Comp Neurol, 363, 642–664.

Carr, D.B. & Sesack, S.R. (2000) GABA-containing neurons in the rat ventral tegmental area project to the prefrontal cortex. Synapse, 38, 114–123.

Casseday, J.H., Kobler, J.B., Isbey, S.F. & Covey, E. (1989) Central acoustic tract in an echolocating bat: an extralemniscal auditory pathway to the thalamus. J Comp Neurol, 287, 247–259.

Constantinidis, C. & Goldman-Rakic, P.S. (2002) Correlated discharges among putative pyramidal neurons and interneurons in the primate prefrontal cortex. Journal of Neurophysiology, 88, 3487–3497.

Corcoran, A.J. & Moss, C.F. (2017) Sensing in a noisy world: lessons from auditory specialists, echolocating bats. J Exp Biol, 220, 4554–4566.

Eiermann, A. & Esser, K.H. (2000) Auditory responses from the frontal cortex in the short-tailed fruit bat Carollia perspicillata. Neuroreport, 11, 421–425.

Esser, K.-H. (2003) Modeling aspects of speech processing in bats––behavioral and neurophysiological studies. Speech Communication, 41, 179–188.

Esser, K.H. & Eiermann, A. (1999) Tonotopic organization and parcellation of auditory cortex in the FM-bat Carollia perspicillata. Eur J Neurosci, 11, 3669–3682.

Feng, A.S. & Ratnam, R. (2000) Neural basis of hearing in real-world situations. Annu Rev Psychol, 51, 699–725.

Funahashi, S., Bruce, C.J. & Goldman-Rakic, P.S. (1993) Dorsolateral prefrontal lesions and oculomotor delayed-response performance: evidence for mnemonic “scotomas”. J Neurosci, 13, 1479–1497.

Fuster, J.M. (2000) Prefrontal neurons in networks of executive memory. Brain Res Bull, 52, 331–336.

Fuster, J.M. (2001) The prefrontal cortex--an update: time is of the essence. Neuron, 30, 319–333.

Garcia-Rosales, F., Beetz, M.J., Cabral-Calderin, Y., Kossl, M. & Hechavarria, J.C. (2018) Neuronal coding of multiscale temporal features in communication sequences within the bat auditory cortex. Commun Biol, 1, 200.

Goldman-Rakic, P.S. (1995) Architecture of the prefrontal cortex and the central executive. Ann N Y Acad Sci, 769, 71–83.

Goldman-Rakic, P.S. & Porrino, L.J. (1985) The primate mediodorsal (MD) nucleus and its projection to the frontal lobe. J Comp Neurol, 242, 535–560.

Goodman, D.F. & Brette, R. (2009) The brian simulator. Front Neurosci, 3, 192–197.

Hackett, T.A., Stepniewska, I. & Kaas, J.H. (1999) Prefrontal connections of the parabelt auditory cortex in macaque monkeys. Brain Res, 817, 45–58.

Hagemann, C., Esser, K.H. & Kossl, M. (2010) Chronotopically organized target-distance map in the auditory cortex of the short-tailed fruit bat. J Neurophysiol, 103, 322–333.

Henny, P. & Jones, B.E. (2008) Projections from basal forebrain to prefrontal cortex comprise cholinergic, GABAergic and glutamatergic inputs to pyramidal cells or interneurons. Eur J Neurosci, 27, 654–670.

Henze, D.A., Borhegyi, Z., Csicsvari, J., Mamiya, A., Harris, K.D. & Buzsaki, G. (2000) Intracellular features predicted by extracellular recordings in the hippocampus in vivo. J Neurophysiol, 84, 390–400.

Kanwal, J.S., Gordon, M., Peng, J.P. & Heinz-Esser, K. (2000) Auditory responses from the frontal cortex in the mustached bat, Pteronotus parnellii. Neuroreport, 11, 367–372.

Kanwal, J.S. & Rauschecker, J.P. (2007) Auditory cortex of bats and primates: managing species-specific calls for social communication. Front Biosci, 12, 4621–4640.

Kayser, C., Logothetis, N.K. & Panzeri, S. (2010) Millisecond encoding precision of auditory cortex neurons. Proc Natl Acad Sci U S A, 107, 16976–16981.

Kobler, J.B., Isbey, S.F. & Casseday, J.H. (1987) Auditory pathways to the frontal cortex of the mustache bat, Pteronotus parnellii. Science, 236, 824–826.

Kossl, M., Hechavarria, J.C., Voss, C., Macias, S., Mora, E.C. & Vater, M. (2014) Neural maps for target range in the auditory cortex of echolocating bats. Curr Opin Neurobiol, 24, 68–75.

Kremer, Y., Leger, J.F., Goodman, D., Brette, R. & Bourdieu, L. (2011) Late emergence of the vibrissa direction selectivity map in the rat barrel cortex. J Neurosci, 31, 10689–10700.

Macias, S., Hechavarria, J.C. & Kossl, M. (2016) Temporal encoding precision of bat auditory neurons tuned to target distance deteriorates on the way to the cortex. J Comp Physiol A Neuroethol Sens Neural Behav Physiol, 202, 195–202.

Martina, M. & Jonas, P. (1997) Functional differences in Na+ channel gating between fast-spiking interneurones and principal neurones of rat hippocampus. J Physiol, 505 (Pt 3), 593–603.

Masquelier, T. (2013) Neural variability, or lack thereof. Front Comput Neurosci, 7, 7.

Medalla, M. & Barbas, H. (2014) Specialized prefrontal “auditory fields”: organization of primate prefrontal-temporal pathways. Front Neurosci, 8, 77.

Mikami, A., Ito, S. & Kubota, K. (1982) Visual response properties of dorsolateral prefrontal neurons during visual fixation task. J Neurophysiol, 47, 593–605.

Murray, E.A. & Rudebeck, P.H. (2018) Specializations for reward-guided decision-making in the primate ventral prefrontal cortex. Nat Rev Neurosci, 19, 404–417.

Newman, J.D. & Lindsley, D.F. (1976) Single unit analysis of auditory processing in squirrel monkey frontal cortex. Exp Brain Res, 25, 169–181.

Ozawa, S., Kamiya, H. & Tsuzuki, K. (1998) Glutamate receptors in the mammalian central nervous system. Prog Neurobiol, 54, 581–618.

Plakke, B., Ng, C.W. & Poremba, A. (2013) Neural correlates of auditory recognition memory in primate lateral prefrontal cortex. Neuroscience, 244, 62–76.

Plakke, B. & Romanski, L.M. (2014) Auditory connections and functions of prefrontal cortex. Front Neurosci, 8, 199.

Puig, M.V., Artigas, F. & Celada, P. (2005) Modulation of the activity of pyramidal neurons in rat prefrontal cortex by raphe stimulation in vivo: involvement of serotonin and GABA. Cereb Cortex, 15, 1–14.

Rao, S.C., Rainer, G. & Miller, E.K. (1997) Integration of what and where in the primate prefrontal cortex. Science, 276, 821–824.

Romanski, L.M., Bates, J.F. & Goldman-Rakic, P.S. (1999a) Auditory belt and parabelt projections to the prefrontal cortex in the rhesus monkey. J Comp Neurol, 403, 141–157.

Romanski, L.M., Tian, B., Fritz, J., Mishkin, M., Goldman-Rakic, P.S. & Rauschecker, J.P. (1999b) Dual streams of auditory afferents target multiple domains in the primate prefrontal cortex. Nat Neurosci, 2, 1131–1136.

Rossant, C., Leijon, S., Magnusson, A.K. & Brette, R. (2011) Sensitivity of noisy neurons to coincident inputs. J Neurosci, 31, 17193–17206.

Sanchez, J.T., Gans, D. & Wenstrup, J.J. (2007) Contribution of NMDA and AMPA receptors to temporal patterning of auditory responses in the inferior colliculus. J Neurosci, 27, 1954–1963.

Stein, R.B., Gossen, E.R. & Jones, K.E. (2005) Neuronal variability: noise or part of the signal? Nat Rev Neurosci, 6, 389–397.

Tanibuchi, I. & Goldman-Rakic, P.S. (2003) Dissociation of spatial-, object-, and sound-coding neurons in the mediodorsal nucleus of the primate thalamus. J Neurophysiol, 89, 1067–1077.

Tomita, H., Ohbayashi, M., Nakahara, K., Hasegawa, I. & Miyashita, Y. (1999) Top-down signal from prefrontal cortex in executive control of memory retrieval. Nature, 401, 699–703.

Uylings, H.B., Groenewegen, H.J. & Kolb, B. (2003) Do rats have a prefrontal cortex? Behav Brain Res, 146, 3–17.

Vogel, A., Hennig, R.M. & Ronacher, B. (2005) Increase of neuronal response variability at higher processing levels as revealed by simultaneous recordings. J Neurophysiol, 93, 3548–3559.

Watanabe, M. (1992) Frontal units of the monkey coding the associative significance of visual and auditory stimuli. Exp Brain Res, 89, 233–247.

Wehr, M. & Zador, A.M. (2003) Balanced inhibition underlies tuning and sharpens spike timing in auditory cortex. Nature, 426, 442–446.

Wilson, F.A.W., Scalaidhe, S.P.O. & Goldmanrakic, P.S. (1994) Functional Synergism between Putative Gamma-Aminobutyrate-Containing Neurons and Pyramidal Neurons in Prefrontal Cortex. P Natl Acad Sci USA, 91, 4009–4013.

Wise, S.P. (2008) Forward frontal fields: phylogeny and fundamental function. Trends Neurosci, 31, 599–608.

Wohlgemuth, M.J., Kothari, N.B. & Moss, C.F. (2016) Action Enhances Acoustic Cues for 3-D Target Localization by Echolocating Bats. PLoS Biol, 14, e1002544.

